# Highly pathogenic avian influenza A H5 virus outbreaks in broiler farms in the Netherlands – Clinical signs, transmission and identification of reporting thresholds

**DOI:** 10.1101/2023.01.05.522008

**Authors:** Jose L. Gonzales, Peter H.F. Hobbelen, Arco N. van der Spek, Edwin P. Vries, Armin R.W. Elbers

**Affiliations:** Wageningen Bioveterinary Research, Lelystad, the Netherlands; Netherlands Food and Consumer Product Safety Authority, Utrecht, the Netherlands

**Keywords:** HPAI, Avian Influenza, Mortality, Broilers, Transmission, Outbreaks

## Abstract

**Background:** For a successful control of highly pathogenic avian influenza virus (HPAIV) epidemics in poultry, early detection is key and it is mostly dependent on the farmer’s prompt identification of disease and reporting suspicions. The latter could be further improved by providing references to farmers for triggering suspicions.

**Methods:** Here we report observations on clinical signs of HPAIV H5N1 and H5N8 infected broiler farms in the Netherlands and analyze their daily mortality and feed and water intake data in order to identify thresholds for reporting suspicions. In addition, mortality data was used to characterize the transmissibility of these viruses, which could help estimate how fast infection spreads within the flock and when infection was likely introduced.

**Results:** The most frequently observed clinical signs in affected flocks were sudden increase in mortality, cyanosis of wattles comb and/or legs and hemorrhagic conjunctiva. Analysis of mortality data indicate that daily mortality higher than 0.17% is an effective threshold for reporting a HPAIV-suspicion. Reliable thresholds for food or water intake could not be stablished. The estimated within-flock transmission rates ranged from 1.1 to 2.0 infections caused by one infectious broiler chicken per day.

**Conclusions:** We identified effective mortality thresholds for reporting suspicions of HPAIV infections. The estimated transmission rates appear to indicate a slow progression of a H5 HPAIV outbreak in affected broiler flocks. The information here provided can be used to improve syndromic surveillance and guide outbreak response.

## 1. Introduction

Highly pathogenic avian influenza A viruses (HPAIV) of the H5 subtypes can infect wild birds and poultry causing severe disease and high mortalities as well as sporadically infect humans (1). Since October 2021, Europe is facing the biggest recorded epidemic caused by HPAI H5 (clade 2.3.4.4b) viruses, predominantly of the H5N1 subtype (2). This epidemic has further spread towards Africa and North America (3). Striking observations during the 2021-2022 epidemic season in Europe are the increased frequency of outbreaks in broiler flocks, compared to previous epidemics (2, 4-9), as well as reports on non-evident clinical signs or absence of increased mortality in affected broiler flocks in Italy (2, 10). The lower frequency of outbreaks in broilers flocks than in other poultry species and production systems observed in previous epidemics has limited the extend at which HPAI infections have been studied in broiler flocks in industrialized production systems. For example, outbreak data from chicken layers, broiler ducks and breeder ducks have been used to stablish reference thresholds for reporting suspicions of HPAI outbreaks based on daily mortality or egg production (11, 12) and similar thresholds have not yet been stablished for broiler flocks. These references can help farmers suspect of potential HPAI outbreaks in their flocks and report this suspicion faster.

In order to improve our understanding of H5 HPAIV infections in broiler flocks, we describe the clinical observations of infected broiler farms in the Netherlands and analyze their data on daily mortality and feed and water intake with the aim of stablishing indicators (thresholds) for reporting suspicions of HPAIV outbreaks, which could help improve syndromic surveillance. Additionally, we use mortality data to quantify the within flock transmission dynamics in infected flocks. This information is relevant to understand how transmissible these H5 HPAIV are in broiler flocks and how fast infection spreads in an infected flock. This information is relevant to guide tracing activities during outbreak response (13, 14).

## 2. Methods

### 2.1. Cases

Observations from six broiler farms infected with H5N1 HPAIV during October-January 2021-2022 and two broiler farms with H5N8 HPAIV during the fall-winter of 2020-2021 were included in the analysis. All eight outbreaks were primary introductions and no transmission towards other farms was observed (3). Data on clinical signs and daily production were collected by the competent authority using standard forms for outbreak response and the production charts of the broiler farms. Seven of these farms had multiple poultry sheds (median 3 (2 – 4)) on the premises. In six of these farms, clinical signs were observed and infection was confirmed in only one poultry shed per farm, and on one farm two out of 4 poultry sheds were infected (Figure 1). Suspicion of infection was detected in the 4^th^-7^th^ week of production, so towards the end of the production period.

**Figure 1.**
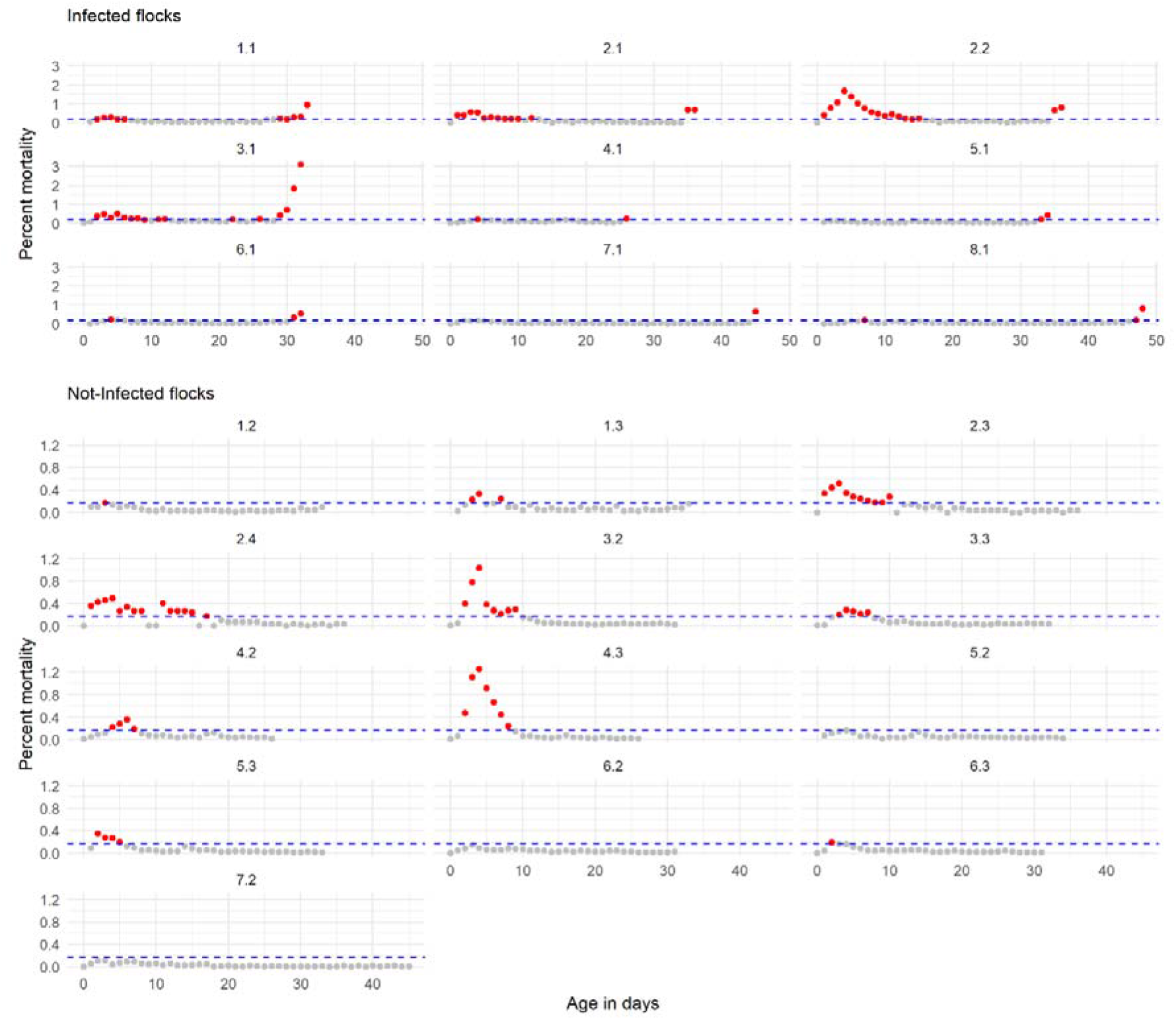
Daily mortality observed in eight broiler farms where outbreaks of HPAIV H5N1 (Farms 1 to 6) and H5N8 (Farms 7 and 8) were confirmed. Farms are identified by number and the decimals represent the flock (shed) identification within each farm. Eg. for Farm 2 the infected flocks are 2.1 and 2.2, and the not infected flocks in this farm are 2.3 and 2.4. The horizontal dashed line indicates the 0.17% mortality threshold. Red dots are days where mortality was higher than this threshold.

### 2.2. Data analysis

#### 2.2.1 Identification of reporting thresholds

##### Daily mortality

A similar analysis approach to that used for broiler ducks (12) for the identification of fixed mortality thresholds were used for this study. Daily mortality rates in “not-infected” flocks (sheds within the farms, n = 13) were modelled using a generalized linear mixed model (GLMM) with a negative binomial distribution (this models fitted better than using a Poisson distribution which showed overdispersion). In this model the daily number of dead broilers was the explanatory variable; the natural log of the daily population size of the flock was the offset, and the age of the broilers in days was the response variable. To account for deviations in linearity in time, natural cubic splines were used on the variable age. The flock identifier nested within the farm identifier were used as the grouping variables (random intercepts, that account for between flock variation on mortality on arrival to the farm (first day)) and age of the broilers as random slope (to account for between flock (within a farm) variation in changes in daily mortality). The best fitting model is provided as supplementary information (Table S1). This model was used to calculate the daily expected mortality and variance (which combines the variance of the fixed and random variables) in a non-infected flock and corresponding 95% upper confidence limits (UCL), estimated by bootstrapping. The latter were considered the maximum mortality rates expected in a non-infected flock and were used to set fixed reporting thresholds. The first (Q1), second (Q2) and third quantiles (Q3) of the estimated UCL were selected and evaluated as reporting thresholds.

##### Feed and water intake

The expected mean and variance of the daily rate of increase in feed (gr of feed/broiler/day) or water intake (ml water/broiler/day) per broiler during the production period were estimated fitting linear mixed models (LMM) to data from the “non infected” flocks. The models that best fitted the data had feed or water intake as the response variable, age in days was the explanatory variable (fixed effect), the farm was introduced as a random intercept (grouping variable) and age in days as random slope (to account for between farm variation in the daily increase in feed or water intake) within each farm. Models that included flock within the farm did not improve the fit, hence were not selected) (Supplementary Tables S2, S3). The estimated variance of increase in feed or water intake was the sum of the fixed and random variance of the expected daily increase in intake. We first assessed the mean expected increase in feed intake - 2.6 times the expected daily standard deviation (SD = square root of variance) as thresholds for the expected maximum drop in feed or water intake. Reduced increase in intake for one or two consecutive days were assessed, but poor specificity was observed (see Results section, Table 2). Additionally we assessed a cero increase or decrease in feed intake, compared with the previous day or two days before, as potential threshold for reporting suspicions.

##### Evaluation of thresholds’ performance

For this evaluation, an “alarm” was defined as an increase in mortality above the mortality threshold or a decrease in feed or water intake bellow the thresholds set by the detection method. A “true alarm” would be an alarm raised any time during a confirmed HPAI outbreak, which was assumed to be up to seven days prior to detection. A “false alarm” would be an alarm raised in non-infected flocks during the production cycle (excluding the first 10 days of age). To evaluate the accuracy of the detection thresholds, we measured the sensitivity (Se), timeliness (T) and the false-positive rate (FP). Se and T were measured on HPAIV-infected flocks and FP was evaluated on non-infected flocks. A detailed description of these parameters is provided in (12).

#### 2.2.2. Quantification of transmission

Daily mortality data was used to quantify the within flock transmission rate β, which is the number of contact infections caused by one infectious bird per day, using a modelling approach and assumption described elsewhere (13).

#### 2.2.3. Data Analysis Software

For the analysis of production parameters and the estimation of both the reporting thresholds and the transmission rate parameter, we used the statistical software package R version 4.0.2 (15). To fit the LMM and GLMM we used the library lme4 (16). Bootstrap estimation of confidence intervals was done using the library merTools (17).

## 3. Results

### 3.1. Clinical signs

Clinical signs observed in the infected flocks (sheds) are summarized in the Table 1. The most frequent observations were sudden increase in mortality, cyanosis of wattles comb and/or legs and hemorrhagic conjunctiva.

**Table 1.**
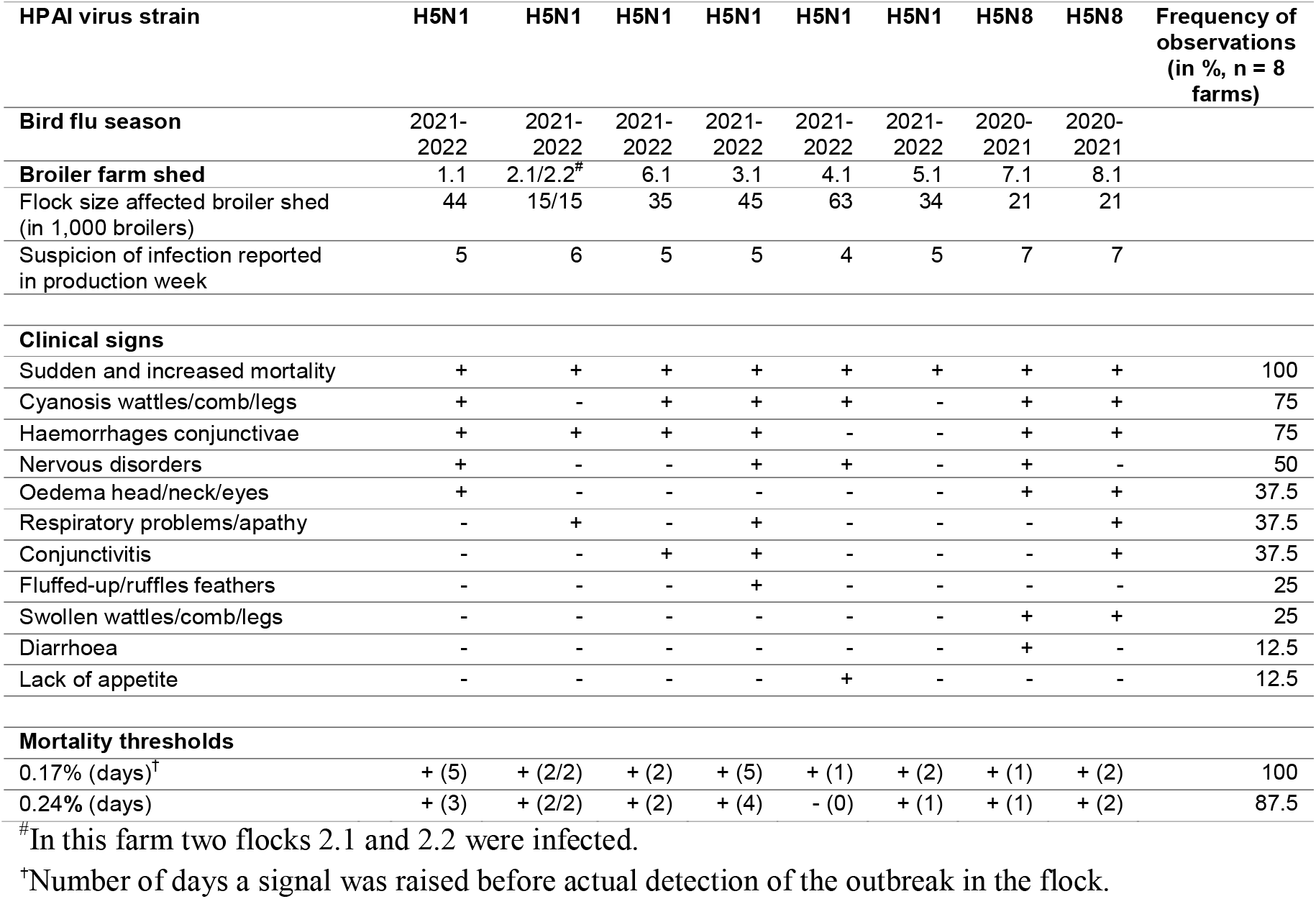
Observed clinical signs in highly pathogenic avian influenza (HPAI) virus H5Nx infected broilers on commercial farms in the Netherlands (2020-2022) and performance of mortality thresholds for raising suspicions in infected flocks.

### 3.2. Establishment of thresholds

Daily mortality in flocks, in the absence of HPAIV infection, was high during the first seven to 10 days of age (Figure 1), which is a common phenomenon in broiler production (18, 19). For the identification of a reporting threshold, we assessed the first and second quantiles of the GLMM predicted 95% upper confidence limits of the daily mortality (age > 10 days). These thresholds were 0.17% and 0.24% daily mortality. Both performed similarly, with the 0.17% threshold being slightly more sensitive and identified one more infected flock in an early stage. Six of the infected farms could have been detected either the same or the previous day the suspicion of infection with HPAIV was reported and two farms could have been detected up to five days earlier (Table 1, Figure 1). The specificity of the thresholds was measured as the number of signals raised per 100 production days (excluding the first 10 days of age) in non-infected flocks (“false alarm” rate (FAR)). The estimated FAR were 2.0 and 1.2 for the 0.17% and 0.24% mortality thresholds, respectively (Table 2). Feed and water intake were also assessed, these thresholds showed high FAR and based on these data we could not establish reliable thresholds for reporting suspicions using these indicators (Table 2). We should however stress the limited number of flocks assessed.

**Table 2.** Identified thresholds and their assessed performance.

| Threshold | Number of detected infected flocks (n = 9) | Number of false alarms per 100 days of production |
| --- | --- | --- |
| Increased daily mortality |  |  |
| 0.17 % | 9 | 2 |
| 0.24 % | 8 | 1.2 |
| Average increases in feed intake per bird |  |  |
| < 3.3 gr (1 day) | 9 | 37.2 |
| < 3.3 gr for 2 consecutive days | 7 | 8.9 |
| <= 0 gr (1 day) | 8 | 24.5 |
| <= 0 gr for 2 consecutive days | 5 | 2.98 |
| <= 0 gr (compared 2 days before) | 6 | 9.76 |
| Average increase in Water intake per bird |  |  |
| < 1.7 ml (1 day) | 9 | 16.2 |
| < 1.7 ml for two consecutive days | 4 | 1.6 |

### 3.3. Transmission parameters

Mortality data suitable for this quantification (exponential increase in mortality observed for more than two consecutive days) was only available for flocks 1.1, 3.1 and 8.1. The estimated transmission rates β ranged from 1.1 to 2 infections per day when assuming a two day latent period and 1.5 to 2 infections per day when assuming a one day latent period (Table 3).

**Table 3.** Transmission parameters estimated based on daily mortality data

| Farm shed id. | Virus strain | Latent period (days)<sup>a</sup> | Infectious period (days)<sup>a</sup> | Transmission rate $\beta$ (95 % CI) (day<sup>-1</sup>) |
| --- | --- | --- | --- | --- |
| 1.1 | H5N1 | 2 | 2.5 | 1.1 (1.0 - 1.3) |
|  |  | 1 | 1.1 | 1.5 (1.4 - 1.7) |
| 3.1 | H5N1 | 2 | 2.5 | 2.0 (1.8 - 2.3) |
|  |  | 1 | 1.1 | 2.0 (1.8 - 2.2) |
| 8.1 | H5N8 | 2 | 2.5 | 1.7 (1.4 - 2.0) |
|  |  | 1 | 1.1 | 1.9 (1.7 - 2.1) |

<sup>a</sup> References and justification for these assumed values are provided by Hobbelen et al (13).

## 4. Discussion

Here we describe the clinical observations made in eight broiler farms affected with H5N8 or H5N1 HPAIV and used the available production and mortality data to identify reporting thresholds as reference for broiler farmers for the identification and reporting of suspicions of HPAIV infections. We observed that in infected Dutch broiler flocks clinical signs and sudden increase in mortality were evident. The most frequent clinical signs observed in affected flocks were sudden increase in mortality, cyanosis of wattles comb and/or legs and hemorrhagic conjunctiva. With the available data we could reliably identify a reporting threshold based on increases in daily mortality and make inferences about the transmissibility of these HPAIV in broiler flocks.

Previous experience has shown that the proportion of broiler chicken flocks affected by HPAIV infection is low (4-9, 20, 21). Assuming that age has no influence in the susceptibility of broilers to HPAIV infection (22), the low frequency of infections in broiler chickens compared to other poultry types could be associated with the short production period. This short production period strongly limits the number of persons, materials and other contacts with the flock. Furthermore, when broilers start production as one-day old chicks, they only need a small volume of fresh air (minimum ventilation) and therefore possible exposure to wind-borne avian influenza virus may be quite limited in the first 2-3 weeks of production. This is coherent with the observation that – as in our present study – often infection in broilers are detected in the last week(s) of production (10, 23, 24).

There are only scarce clinical observations on field outbreaks in broiler farms. During the large HPAI H7N7 epidemic in the Netherlands in 2003, three broiler farms were infected out of a total 240 infected poultry – predominantly egg-producing – farms (8). The most striking clinical signs observed in the broiler farms were sudden and rapidly increasing mortality and respiratory disorders. In Cambodia, typical clinical features in broilers after HPAI H5N1 infection were sudden death, severe depression, edema of neck and wattles, swollen and cyanotic combs and wattles and congested conjunctiva (24). An exceptional situation enrolled in the poultry-dense area of Veneto, in the North-East of Italy (10). Turkey and broiler farms were the most affected poultry species. Special monitoring and control activities – sampling of dead broilers during the production period - indicated that several broiler flocks were infected in the absence of increased mortality and/or clinical signs. Most infected broiler flocks were detected infected just prior to shipment of the broilers to the slaughterhouse. Except for the Italian experience, the clinical observations in our present study correlate with the scarce other information available. The absence of observed signs in Italy could also be a result of detection before sudden or exponential increase in mortality started as reported by following one of the infected flocks (10). This early detection in Italy was a result of the implementation of the special monitoring measures mentioned above.

An effective reporting threshold should be one which leads to early detection and rapid control of infection (benefit) whilst minimizes the costs associated with rapid responses to false alarms. Following this principle and with the limited number of farms available for analysis, we could only identify mortality thresholds as effective indicators to trigger suspicions. Daily mortality higher than 0.17% led to the sensitive detection of all outbreaks in this study whilst limiting false alarms to ≤1 per production cycle. This threshold aligns with observations done in early detected infected flocks in Italy where mortalities lower than 0.2% were reported (10). When comparing with other production systems the 0.17% threshold is higher than that identified for chicken layers (0.13%) (11) and lower than the threshold for broiler ducks (0.3%) (12). Although we use a limited number of farms, we consider these estimated threshold as valuable. By using this threshold, we see that detection of HPAIV-infection in some flocks could have been reported up to five days earlier.

For three of the outbreaks there was enough information to quantify transmission. Irrespective of the assumptions made on the latent and infectious period, we estimated β ranging from 1.1 to 2.0 day^-1^ for both the H5N1 and the H5N8 HPAIV. These estimates are similar to those estimated for H5N8 HPAIV infected broiler flocks (n = 12) between 2020-2021 in Japan. The median (range) β was 1.45 (0.66 – 3.39) day^-1^ (25). Taken these studies together, they appear to indicated that the rate of transmission of these H5 HPAIV in broiler flocks appears to be slower than in layer flocks (β > 4.4 day^-1^) (13) and as a consequence the daily increase in mortality will be lower in broilers than layers as it was also observed during the H7N7 HPAI epidemic in the Netherlands (26). Therefore, assuming that the case fatality is similar between broilers and layers, one could expect that the time taken to observe increased mortality in broilers and thereby detection would be longer than that expected for layer farms. This could explain the observations in Italy, where increased mortality was only observed eight days later than laboratory detection following emergency (active) surveillance activities (10). Hence, tracing back activities following infections in broilers should consider a longer introduction window than layers.

We should acknowledge the limitations of this study due to the small number of farms assessed. This small number of observations may have limited for example the capacity to identify other thresholds such as feed and water intake as well as the evaluation, with higher confidence, of the performance of the mortality threshold here suggested. Validation of the performance of this and previously suggested thresholds for other poultry production systems is required to increase the confidence in their efficacy and stimulate their application.

Finally, it should be noted that farmers, whose farms were included in this study, reported suspicions early in the within-flock infection process. In some cases they reported the first day they observed an unusual increase in mortality, even though this increase was lower than the threshold for reporting suspicions that is now applicable (mortality > 0.5%) (27). This early reporting and detection may have contributed to a reduction of the risk of transmission to other farms. These observations highlight the importance of early detection, which can be further optimized by providing farmers with effective reporting thresholds, to guide them in reporting suspicions.

## Acknowledgments

This research was supported by The Netherlands’ Ministry of Agriculture, Nature and Food Quality (Statutory Task Unit Infectious Animal diseases: WOT-01-001-004).

## Authors’ Contributions

Conception and design of the study: Gonzales JL and Elbers RW; performed data analysis and interpretation: Gonzales JL, Hobbelen P and Elbers RW; performed data acquisition van der Spek AN and Vries EP.

## Availability of Data and Materials

All data used for this analysis is provided as supplementary material.

## Ethical Approval and Consent to Participate

Not applicable

## Consent for Publication

Not applicable

