## Supplementary material for "Highly pathogenic avian influenza A H5 virus outbreaks in broiler farms in the Netherlands – Clinical signs, transmission and identification of reporting thresholds": Table S1, S2, S3

*Mortality*

Table S1. Negative binomial GLMM fitted to model daily mortality in not infected broiler flocks. Parameter values are presented as log estimates.

| Knots for natural spline effects with 2 degrees of freedom | | | | |
| --- | --- | --- | --- | --- |
| Knot number | Age |  |  |  |
| 2 | 22 |  |  |  |
| Variable | Estimate | std. error | Z value | p.value |
| Intercept (Arrival age) | -7.23 | 0.18 | -40.56 | <0.001 |
| age, ns 1 | -1.69 | 0.39 | -4.31 | <0.001 |
| age, ns 2 | 0.17 | 0.41 | 0.41 | 0.68 |
| *Random Effects* |  |  |  |  |
| Farm:flock^1^ |  | 0.84 |  |  |
| Farm:flock:age^2^ |  | 0.04 |  |  |
| Correlation house and flock |  | -0.93 |  |  |
| Farm^3^ |  | 0.23 |  |  |
| Observations | 297 (6 farms, 6 flocks) | |  |  |
| Conditional R2 | 0.83 |  |  |  |

^1^Between flock nested within the farm variation at day (age) of arrival to farm.

^2^Between flock variation in changes in daily mortality

^3^Between farm variation at arrival day

*Feed and water intake*

Table S2. Linear mixed (best fit) model fitted to the daily feed intake data from not-infected broiler flocks.

| Variable | Estimate | std.error | Z value | df | p.value |
| --- | --- | --- | --- | --- | --- |
| Intercept (Arrival age) | 3.25 | 2.75 | 1.18 | 5.78 | 0.284483 |
| Age | 5.12 | 0.24 | 21.39 | 5.84 | 0.000001 |
| *Random variables* |  |  |  |  |  |
| Farm^1^ |  | 6.81 |  |  |  |
| Farm:age^2^ |  | 0.62 |  |  |  |
| Residual SD |  | 9.49 |  |  |  |
| Correlation Farm and age |  | 0.66 |  |  |  |
| Observations | 416 |  |  |  |  |
| Conditional R2 | 0.97 |  |  |  |  |

^1^ Between farm variation in mortality at baseline (arrival age/day)

^2^ Between farm variation in the rate of increase in feed intake

Table S3. Linear mixed (best fit) model fitted to the daily water intake data from not-infected broiler flocks.

| Variable | Estimate | std.error | Z value | df | p.value |
| --- | --- | --- | --- | --- | --- |
| Intercept (Arrival age) | 12.12 | 3.08 | 3.93 | 6.00 | 0.01 |
| Age | 7.65 | 0.82 | 9.31 | 6.01 | 0.00 |
| *Random effects* |  |  |  |  |  |
| Farm^1^ |  | 7.61 |  |  |  |
| Farm:age^2^ |  | 2.17 |  |  |  |
| Residual SD |  | -0.42 |  |  |  |
| Correlation Farm and age |  | 11.17 |  |  |  |
| Observations | 426 |  |  |  |  |
| Conditional R2 | 0.98 |  |  |  |  |

^1^ Between farm variation in mortality at baseline (arrival age/day)

^2^ Between farm variation in the rate of increase in feed intake
